# Diversity and evolution of cerebellar folding in mammals

**DOI:** 10.1101/2022.12.30.522292

**Authors:** Katja Heuer, Nicolas Traut, Alexandra A. de Sousa, Sofie Valk, Roberto Toro

## Abstract

The process of brain folding is thought to play an important role in the development and organisation of the cerebrum and the cerebellum. The study of cerebellar folding is challenging due to the small size and abundance of its folia. In consequence, little is known about its anatomical diversity and evolution. We constituted an open collection of histological data from 56 mammalian species and manually segmented the cerebrum and the cerebellum. We developed methods to measure the geometry of cerebellar folia and to estimate the thickness of the molecular layer. We used phylogenetic comparative methods to study the diversity and evolution of cerebellar folding and its relationship with the anatomy of the cerebrum. Our results show that the evolution of cerebellar and cerebral anatomy follows a stabilising selection process. Ancestral estimations indicate that size and folding of the cerebrum and cerebellum increase and decrease concertedly through evolution. Our analyses confirm the strong correlation between cerebral and cerebellar volumes across species, and show that large cerebella are disproportionately more folded than smaller ones. Compared with the extreme variations in cerebellar surface area, folial wavelength and molecular layer thickness varied only slightly, showing a much smaller increase in the larger cerebella. These findings provide new insights into the diversity and evolution of cerebellar folding and its potential influence on brain organisation across species.

## Introduction

Brain folding may play an important role in facilitating and inducing a variety of anatomical and functional organisation patterns (Welker 1990, Toro 2012). Among the many folded structures of the mammalian brain, the cortices of the cerebrum and the cerebellum are the two largest ones. Both cortices are organised in layers, with neuronal cell bodies occupying a superficial position and principal neurones – pyramidal neurones in the cerebral cortex, Purkinje cells in the cerebellum – sending axons towards an internal white matter. Many aspects of the organisation, development and evolution of the cerebral and cerebellar cortex are, however, very different. It is interesting then to compare folding in both structures as a first step towards understanding how the mechanics of folding could influence the organisation in these two different types of tissue (Franze 2013, Kroenke and Bayly 2018, Foubet et al. 2019, Heuer and Toro 2019).

While the cerebral cortex is unique to mammals, the cerebellum is present in all vertebrates, with cerebellum-like structures which can be even identified in invertebrates such as octopuses (Shigeno et al. 2018). While the main target of pyramidal cells is the cerebral cortex itself (through profuse cortico-cortical connections), Purkinje cell afferents are mostly feedforward. While dendritic trees of pyramidal neurones can be found in all cortical layers, those of Purkinje cells are restricted to the molecular layer – their nuclei forming a boundary between the molecular and granular layers.

The process leading to folding in both the cerebrum and the cerebellum is likely to be a mechanical instability (buckling) triggered by the growth of the external layers on top of the elastic core (Toro and Burnod 2005, Tallinen et al. 2014, Foubet et al. 2019, Heuer and Toro 2019, Lawton et al. 2019, Llinares-Benadero and Borrell 2019, Van Essen 2020). However, while the folds of the cerebral cortex seem to be able to take any orientation (especially in large gyrencephalic brains such as those of humans or cetaceans), the cerebellar folia are almost completely parallel to one another, accordion-like, forming the same characteristic type of tree-like structures from mice, to humans and whales (Leto et al. 2016).

The different folding patterns may reflect the different nature of cortical expansion in both structures. In both, the growth of neuronal dendritic trees may be a main source for the increase in cortical volume. In the case of the cerebrum, the dendritic trees of pyramidal neurones grow to occupy a 3-D volume (Jiang et al. 2020), leading to the homogeneous expansion of the cortical surface and the formation of folds with a variety of orientations. In the case of the cerebellum, however, the growth of the dendritic trees of Purkinje cells is largely restricted to a 2-D plane (Kaneko et al. 2011), leading to dendritic trees which are parallel to one another, and orthogonal to the main direction of folding.

Here we aimed at characterising the folding of the cerebellar cortex using histological data from a sample of 56 mammalian species. The analysis of cerebellar folding is made difficult by the small size of the folia. Magnetic resonance imaging, the main tool for studying the folding of the cerebral cortex, does not provide the resolution required to distinguish and reconstruct cerebellar folding. Histological data, even at low scanning resolution, can provide sufficient information to distinguish folia as well as cerebellar cortical layers. However, data is only 2-D, and sectioning often induces non-trivial deformations which make it challenging to create full 3-D reconstructions (see, however, the work of Sereno et al. 2020 and Zheng et al. 2022). Using histological data Ashwell (2020) was able to estimate the surface area in 90 species of marsupials and 57 species of eutherian mammals, but did not study folia, of which the measurement remains challenging.

In our analyses we manually segmented the surface of the cerebellum and developed a method to automatically detect the boundary between the molecular and granular layers, allowing us to estimate the median thickness of the molecular layer. We also developed a method to detect individual folia, and to measure their median width and perimeter. Buckling models of brain folding suggest an inverse relationship between folding frequency and cortical thickness, with thicker cortices leading to lower frequency folding (Toro and Burnod 2005). This has been verified in the mammalian cerebrum, and we aimed at studying it in the cerebellum (Mota and Herculano-Houzel 2015). We used phylogenetic comparative methods to study the evolution of the cerebellum and its relationship with the cerebrum and with body size.

Our results confirm the strong allometry between the size of the cerebellum and the cerebrum across mammals (Herculano-Houzel 2010, Smaers et al. 2018, Ashwell 2020) and show similarly strong relationships for the width of cerebellar folds and the thickness of the molecular layer. This deeply conserved pattern suggests the presence of a common mechanism underlying the development of the cerebellum and cerebellar folding across mammals. All our data and code are openly accessible online.

## Materials and methods

### Histological data

Most of the data used in this study comes from the Brain Museum website (https://brainmuseum.org) – the Comparative Mammalian Brain Collection initiated by Wally Welker, John Irwin Johnson and Adrianne Noe from the University of Wisconsin, Michigan State University and the National Museum of Health and Medicine. This is the same eutherian mammal data used by Ashwell (2020) through the Neuroscience Library website (https://neurosciencelibrary.org, offline since June 2022). The human brain data comes from the BigBrain Project (https://bigbrainproject.org) led by teams from Forschungszentrum Jülich and McGill University (Amunts et al. 2013). The rhesus macaque data is part of the BrainMaps project (http://brainmaps.org) by Edward G. Jones at UC Davis. (Fig. 1 and Supplemental Table S1 for a detailed list).

**Figure 1.**
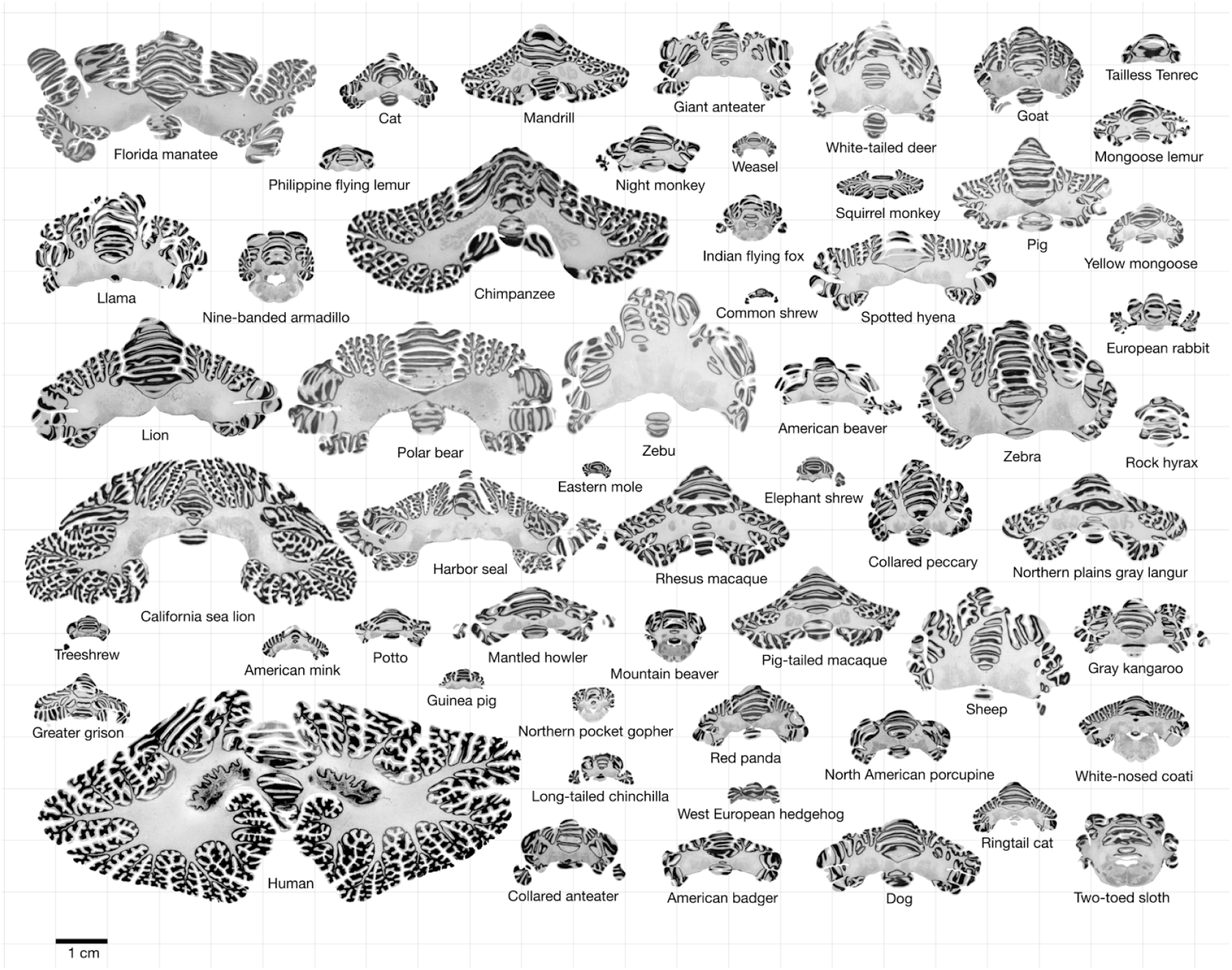
Data. All mid-cerebellar slices for the different species analysed, at the same scale. This dataset is available online for interactive visualisation and annotation: https://microdraw.pasteur.fr/proiect/brainmuseum-cb.

We used our Web tool MicroDraw to visualise and segment the histological data online (https://microdraw.pasteur.fr). We downloaded the Brain Museum data, and used the vips library (https://www.libvips.org) to convert it to deep zoom image format – the multi-scale image format required by MicroDraw. The data from the BigBrain and BrainMaps projects is already encoded in multi-scale format, and we only wrote translation functions to allow access through MicroDraw. The Brain Museum Cerebellum project can be accessed at https://microdraw.pasteur.fr/project/brainmuseum-cb.

The resolution of the histological images in the Brain Museum was very variable. We considered that image resolution was in all cases sufficient to estimate the length and area of the cerebellum sections, however, we excluded some species from our estimation of molecular thickness when resolution was deemed insufficient. For this, we plotted the total number of pixels of the image versus cerebellum size, and excluded species where the density of pixels per mm^2^ was lower than 3.5 – a threshold determined by visual inspection. The datasets from the BrainMaps (rhesus macaque), and the BigBrain project (human) were all high resolution (1-20μm).

### Segmentation

We used the polygon tool in MicroDraw, with different colours to draw the contour of the mid-slice of the cerebellum, the mid-slice of the cerebrum and the scale bar. We used a Google spreadsheet to keep track of species information, and to identify slices of interest and coordinate our work. In the case of the human brain from the BigBrain project, and the rhesus macaque from the BrainMaps project, where no scale bar is present directly in the slices, we obtained scale information from the websites.

### Measurement of section area and length of the cerebral and cerebellar cortex

Table 1 summarises the variables used in our analyses and their sources. We used the python package microdraw.py (https://github.com/neuroanatomy/microdraw.py) to query MicroDraw’s API and programmatically download all vectorial segmentations and images (Fig. 2). We computed the length of the cerebral and cerebellar segmentation contours, and used the scale bar information to convert the data to millimetres. We used an artificial calibration image to ensure that our algorithms produce the correct measurements. Measurements were not modified to account for shrinkage, and reflect the raw data obtained from the images (see Ashwell (2020) for a discussion of shrinkage in the Brain Museum collection).

**Figure 2.**
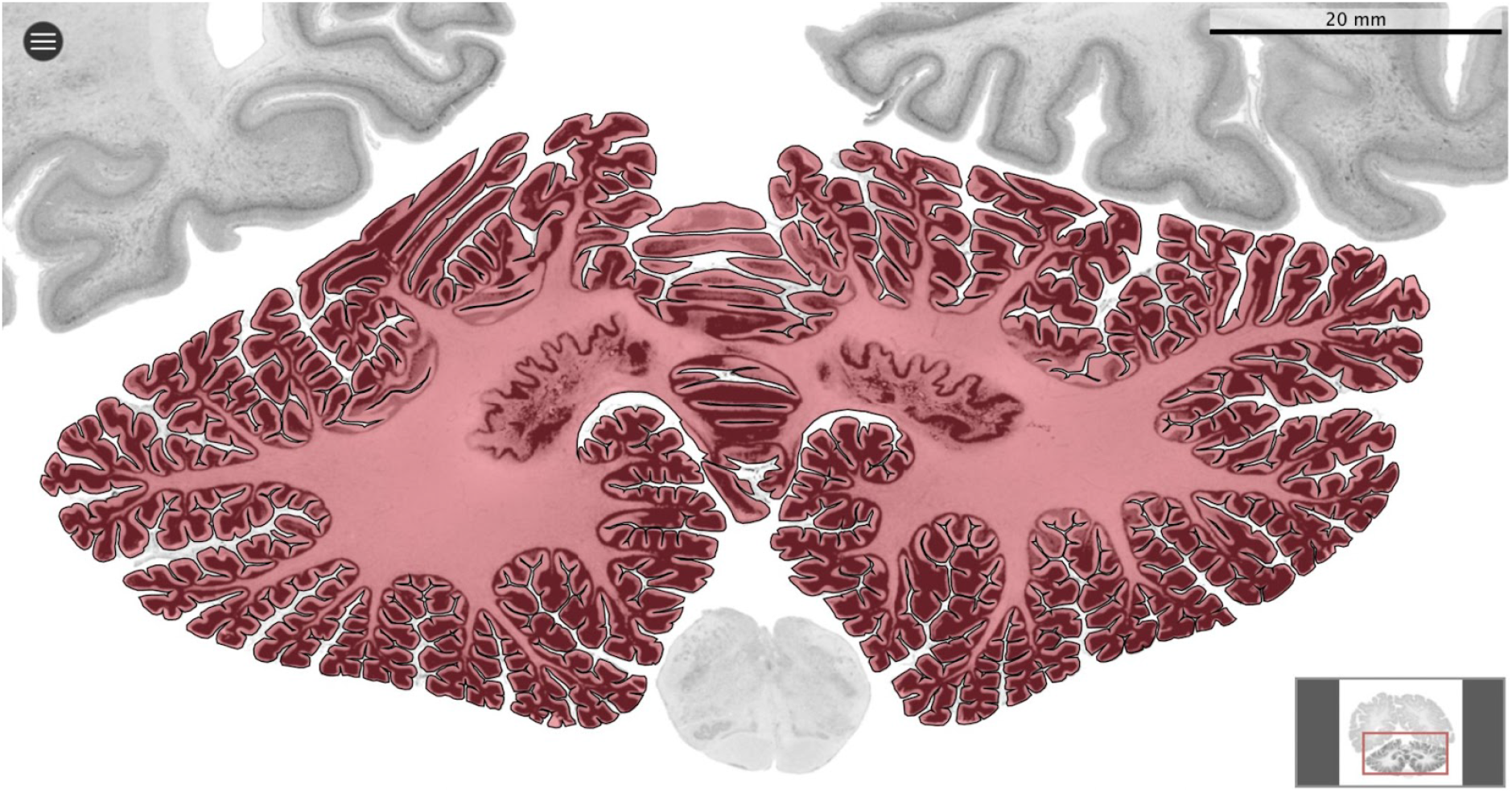
Segmentation. All datasets were indexed in our collaborative Web application MicroDraw (https://microdraw.pasteur.fr) to interactively view and annotate the slices. The contour of the cerebellum was drawn manually using MicroDraw’s drawing and boolean operation tools. The example image shows the human cerebellum from the BigBrain 20μm dataset (Amunts et al. 2013).

**Table 1.** Definition of the variables used in the current study and their sources.

| Variable name | Definition | Source |
| --- | --- | --- |
| Cerebellar section area | Area of the cerebellar mid-slice section (mm<sup>2</sup>; Fig. 2) | Measured |
| Cerebellar section length | Perimeter around the cerebellar pial surface of mid-slice section (mm; Fig. 2) | Measured |
| Folial width (cerebellum) | Euclidean distance between the two flanking sulci of a folium, averaged for all folia in the mid-slice section (mm; Fig. 3b) | Measured |
| Folial perimeter (cerebellum) | Perimeter along the cerebellar pial surface between the two flanking sulci of a folium, averaged for all folia in the mid-slice section (mm; Fig. 3c) | Measured |
| Thickness (cerebellum) | Thickness of the molecular layer for the cerebellar mid-slice section, averaged from the lengths profile lines that bisect the molecular layer from its border with the pial surface to its border with the granular surface (mm; Fig. 4) | Measured |
| Cerebral section area | Area of the cerebral mid-slice section (mm<sup>2</sup>) | Measured |
| Cerebral section length | Perimeter around the cerebral pial surface of mid-slice section (mm) | Measured |
| Brain weight | Weight of the whole brain, including cerebrum and cerebellum (g) | Ballarin et al. 2016, Burger et al. 2019, Smaers et al. 2021 |
| Body weight | Weight of the whole body (g) | Ballarin et al. 2016, Burger et al. 2019, Smaers et al. 2021 |
| Cerebellar volume | Volume of the entire cerebellum, estimated from 2-D histological sections or MRI (mm<sup>3</sup>) | Smaers et al. 2018 |
| Cerebral volume | Volume of the entire cerebrum, estimated from 2D histological sections or MRI (mm<sup>3</sup>) | Smaers et al. 2018 |
| Sultan's folial width | Length of a cerebellar fold in a cross-section orthogonal to the fold axis: each fold may contain several individual folia. | Sultan and Braitenberg 1993 |

### Measurement of folial width and perimeter

We resampled the manual segmentations to have a homogeneous density of vertices and automatically detected sulci (Fig. 3). At each vertex we measured the mean curvature of the segmentation path. We smoothed the mean curvature measurements at 10 different scales, which produced for each vertex a 10-D signature. We used a random forest algorithm to automatically distinguish 3 classes of vertices: gyri, sulci and walls. We measured the width of each folium as the Euclidean distance between its 2 flanking sulci. We measured the perimeter of each folium as the distance along the cerebellar surface between its two flanking sulci. For each species we used the median width and the median perimeter across all folia, to make our estimations robust to outliers.

**Figure 3.**
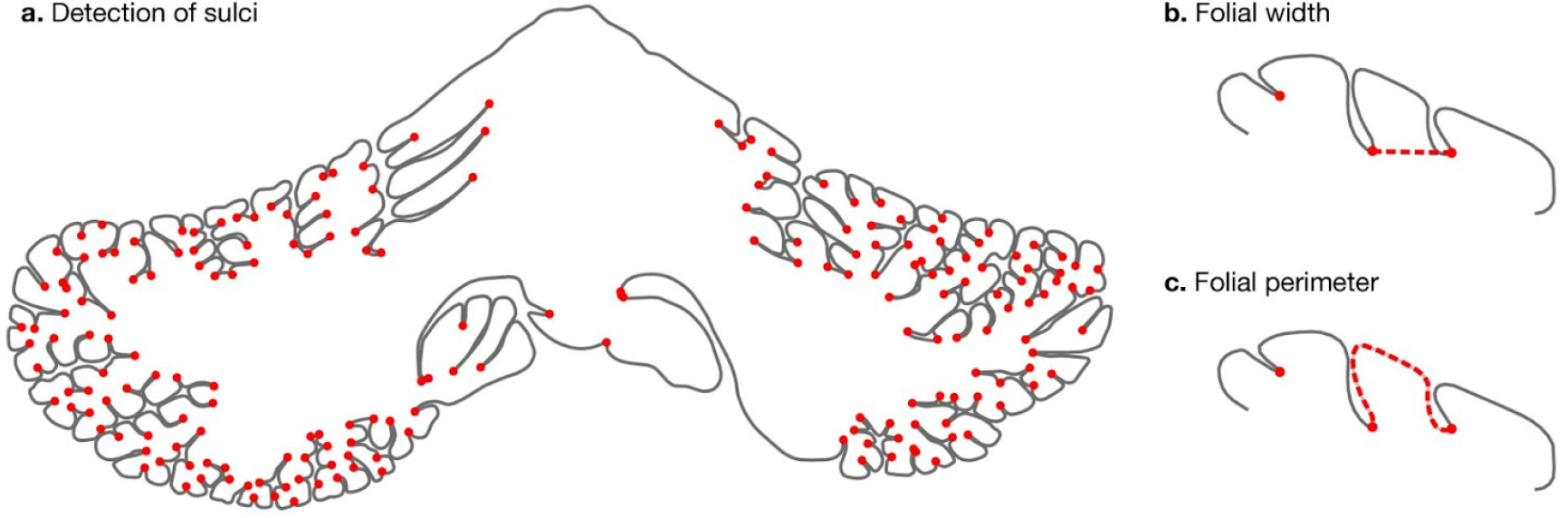
Detection of sulci and measurement of folial width and perimeter. Sulci were automatically detected using mean-curvature filtered at different scales **(a)**. From this, we computed folial width **(b)** and folial perimeter **(c)**.

### Measurement of molecular layer thickness

We estimated the thickness of the molecular layer automatically (Fig. 4). We processed the histology images to convert them to grey levels, denoise them and equalise their histograms, using functions from the Scikit-image package (van der Walt et al. 2014). Starting from the manual segmentation, we created a binary mask and masked out the non-cerebellar regions. In mature cerebella, the molecular layer appears as a light band followed by the granular layer which appears as a dark band. We applied a Laplacian smoothing to the masked image, which created a gradient going from white close to the surface of the cerebellum to black toward the granular layer. We computed the gradient of the image, which produced a vector field from the outside to the inside. We integrated this vector field to produce a series of scan lines, each for every vertex in the cerebellar segmentation. The vector field was integrated only if the grey values decreased, and if a minimum was reached, the vector field was continued linearly. At the end of this procedure, all scan lines had the same length, computed so as to cover the whole range from the surface of the cerebellum to the granular layer. The grey levels in the original image were sampled along each scan line, producing a grey level profile. The grey level profiles were derived, and a peak detection function was used to determine the point of maximum white/grey contrast, indicating the boundary between the molecular and granular layers. For each scan line the corresponding thickness of the molecular layer was defined as the length from the surface until the maximum contrast point. Finally, for each species, the median thickness was computed, to render our estimation robust to outliers.

**Figure 4.**
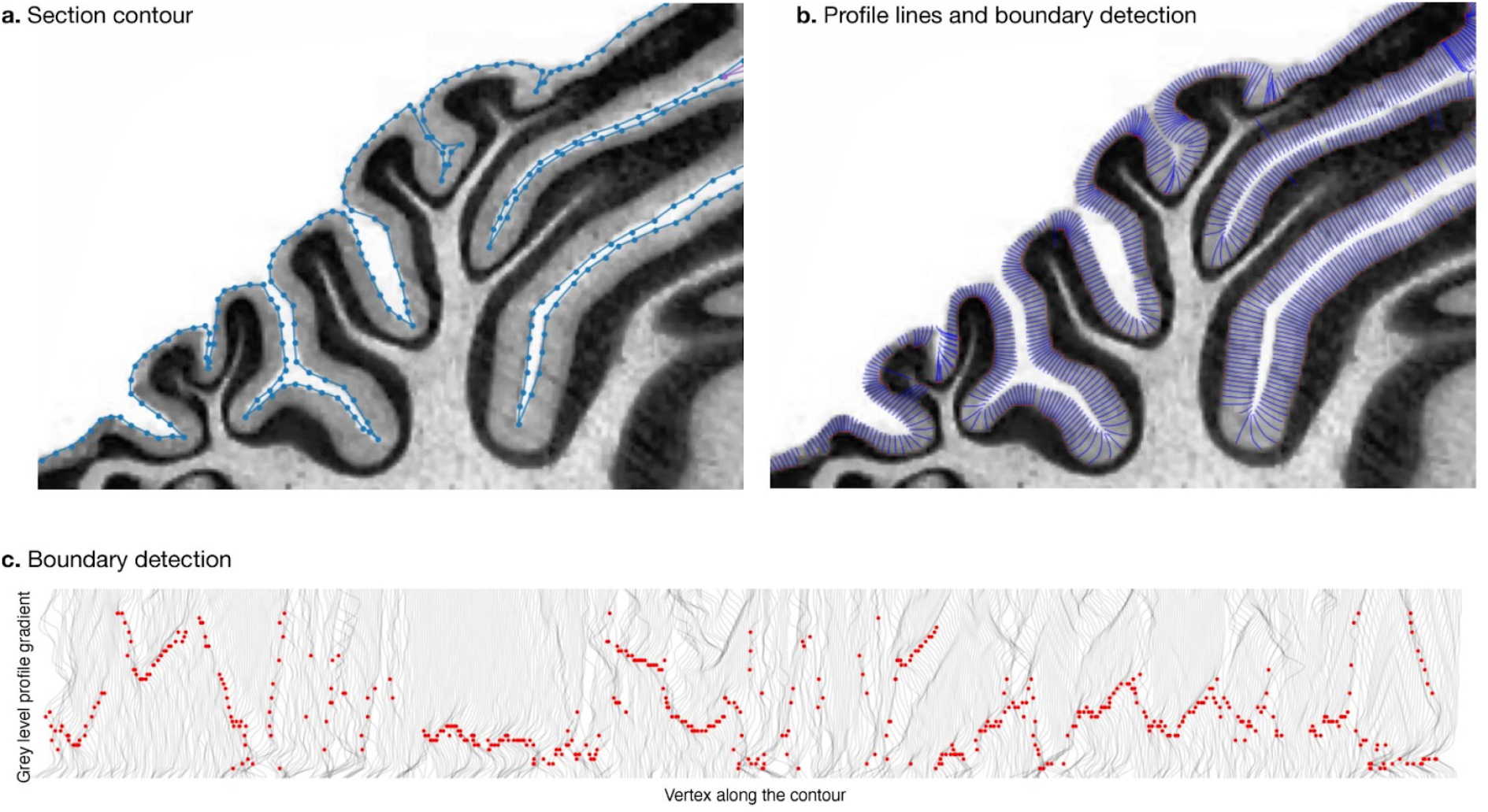
Measurement of molecular layer thickness. The thickness of the molecular layer of the cerebellum was measured automatically using the manual segmentation of the surface. **a.** Manually annotated surface contour. **b.** Automatically computed profile lines. **c.** Profile grey level and border detection. Red dots: detected borders.

### Phylogenetic comparative analyses

We used phylogenetic least squares and phylogenetic principal components to look at bivariate and multivariate relationships among neuroanatomical phenotypes with the mvMorph package (Clavel et al. 2015). We obtained a phylogenetic tree for our 56 species from the TimeTree website (Fig. 5, https://www.timetree.org, Kumar et al. 2017). Phylogenetic comparative methods use this tree to determine an expected correlation structure for the data.

**Figure 5.**
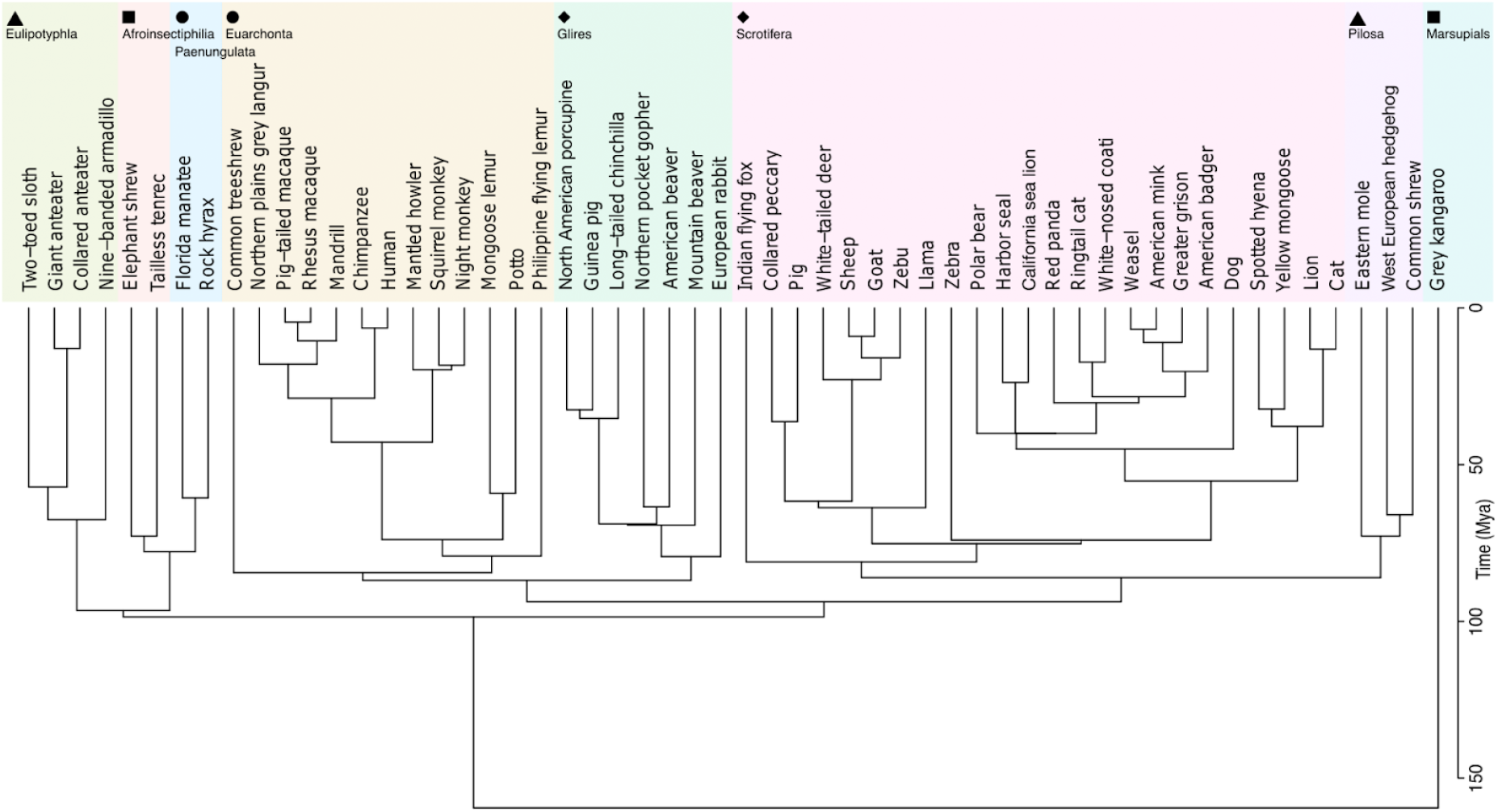
Phylogenetic tree. The phylogenetic tree for 56 species was downloaded from the TimeTree website (http://www.timetree.org, Kumar et al. 2017) and coloured in 8 groups based on hierarchical clustering of the tree.

We estimated ancestral values from the tip species of the phylogenetic tree to the common ancestor using different evolutionary models: the Brownian Motion model (BM) which assumes that phenotypes vary randomly across time, the Ornstein-Uhlenbeck model (OU) which assumes in addition the presence of an optimal value for each phenotype (stabilising selection), and the Early Burst model (EB), which assumes that the speed of phenotypic variation may be different across time. The best fitting model was selected by comparing the Generalised Information Criterion. We used this model to impute missing data, and to generate an estimation of ancestral phenotypes. Phylogenetic comparative analyses used the R packages phytools (Revell 2011) and mvMorph (Clavel et al. 2015). The code necessary to reproduce our statistical analyses is available at https://github.com/neuroanatomy/comp-cb-folding.

## Results

### Validation of measurements: Cerebellar and cerebral section areas are a good proxy for total volume

Our measurements correlated well with those reported in the literature. We segmented the coronal mid-slice of the cerebellum and the cerebrum. These section areas should scale as the 2/3 power of the volume. We confirmed that this was the case by comparing our measurements with the volume measurements reported by Smaers et al. (2018): the correlation captured ~96% of the variance (Fig. 6a, b). Additionally, we observed a strong correlation between cerebellum section area and the volume of the medial cerebellum (although we do not use medial cerebellar measurements in our analyses). See Supplemental Fig. 1 for correlation with measurements in Ashwell (2020).

**Figure 6.**
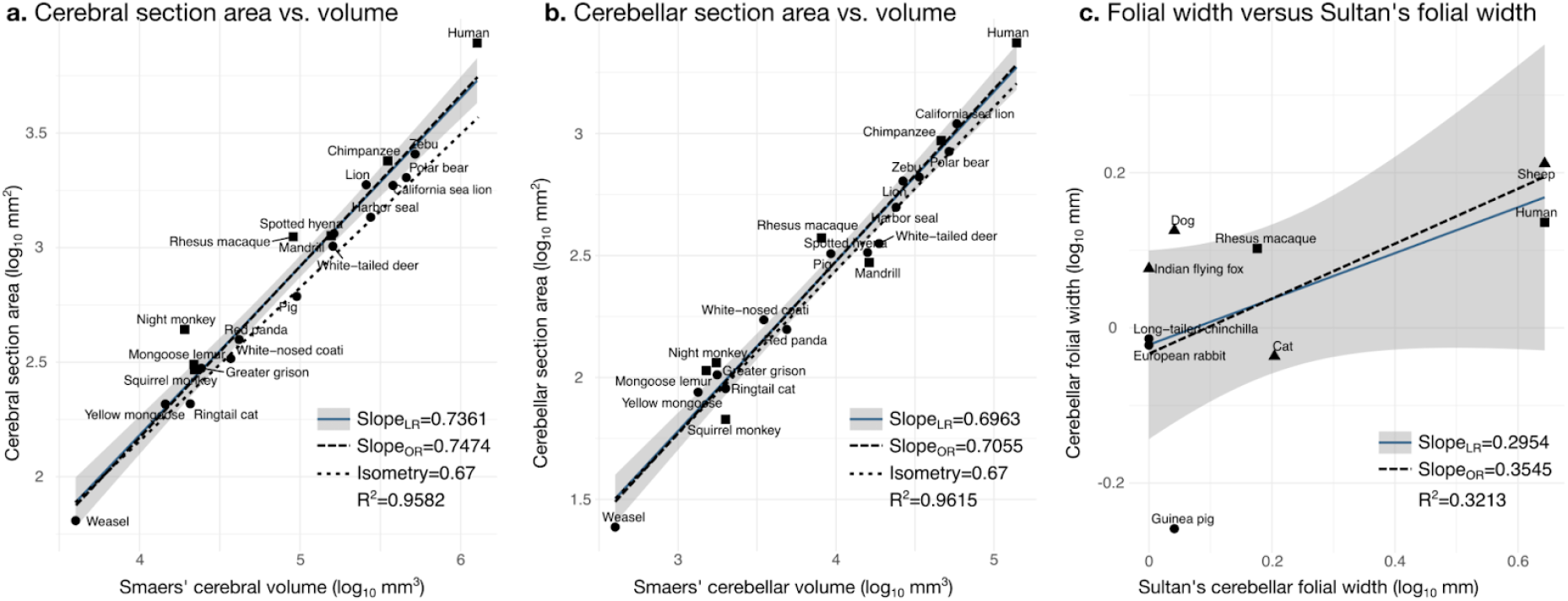
Validation of neuroanatomical measurements. Comparison of our cortical section area measurement **(a)** and cerebellar section area **(b)** with volume measurements from Smaers et al. (2018). **(c)** Comparison of our folial width measurement with folial width from Sultan and Braitenberg (1993). The folial width measurement reported by Sultan and Braitenberg, which may include several folia, do not correlate significantly with our measurements (2-tailed p=0.112). LR: Linear regression with 95% confidence interval in grey. OR: Orthogonal regression.

There are no comparative analyses of the frequency of cerebellar folding, to our knowledge. The closest is the measurements reported by Sultan and Braitenberg (1993), measuring the length of structures which may include several individual folia. Our measurements of cerebellar folial width and folial perimeter, however, do not correlate significantly with Sultan and Braitenberg’s measurements (Fig. 6c). Ashwell (2020) studied cerebellar folding through a foliation index, defined as the ratio of the cerebellar pial surface over the external cerebellar surface. This method, however, does not provide information about folding frequency, and would be unable to distinguish a cerebellum with many shallow folds from one with a few deep ones, which could both produce a similar foliation index.

### The evolution of cerebellar and cerebral neuroanatomy follows a stabilising selection process

We compared 3 different approaches to model the evolution of cerebellar and cerebral neuroanatomical measurements: Brownian Motion, Ornstein-Uhlenbeck (stabilising selection) and Early-Burst. Comparing the goodness of fit of for the 3 models we observed substantial evidence in favour of the Ornstein-Uhlenbeck model (Table 2, smaller values indicate a better fit). The second most likely model was Brownian Motion.

**Table 2.**
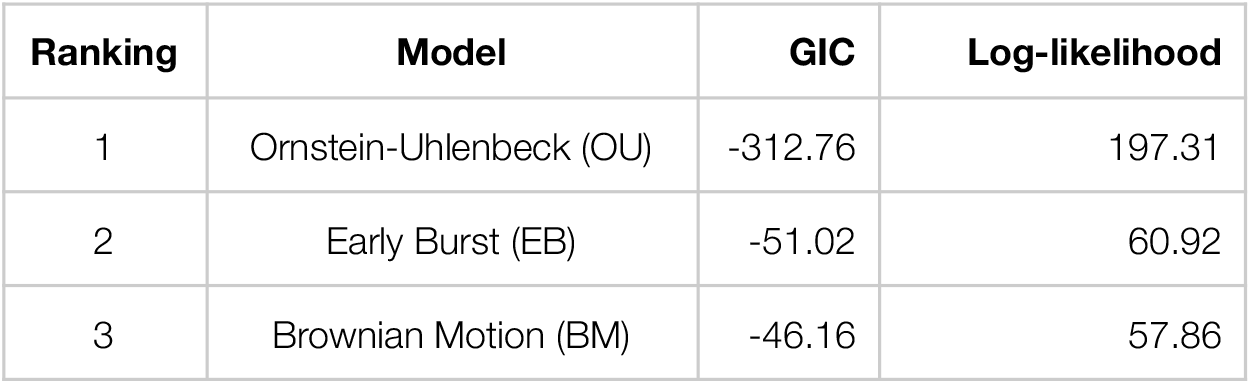
Ranking of phylogenetic comparative model fit based on the Generalised information criterion (GIC). The best fit to the data (smallest GIC value) was obtained for the Ornstein-Uhlenbeck model of stabilising selection.

| Ranking | Model | GIC | Log-likelihood |
| --- | --- | --- | --- |
| 1 | Ornstein-Uhlenbeck (OU) | -312.76 | 197.31 |
| 2 | Early Burst (EB) | -51.02 | 60.92 |
| 3 | Brownian Motion (BM) | -46.16 | 57.86 |

The analyses of relationships among phenotypes that follow use the OU model to control for phylogenetic structure (otherwise, the observation would not be evolutionarily independent).

### Correlation structure: Cerebellar size and folding correlate strongly with cerebral size and folding

All measurements were strongly correlated, especially cerebellar and cerebral section area, surface contour length, total brain weight and total body weight (Fig. 7a, analyses included phylogenetic structure). To better understand the relationships among variables we computed partial correlations, which provide a more local view on the relationships among pairs of measurements conditional to all remaining measurements. We also performed a phylogenetic principal component analysis to study the main patterns of neuroanatomical diversity.

**Figure 7.**
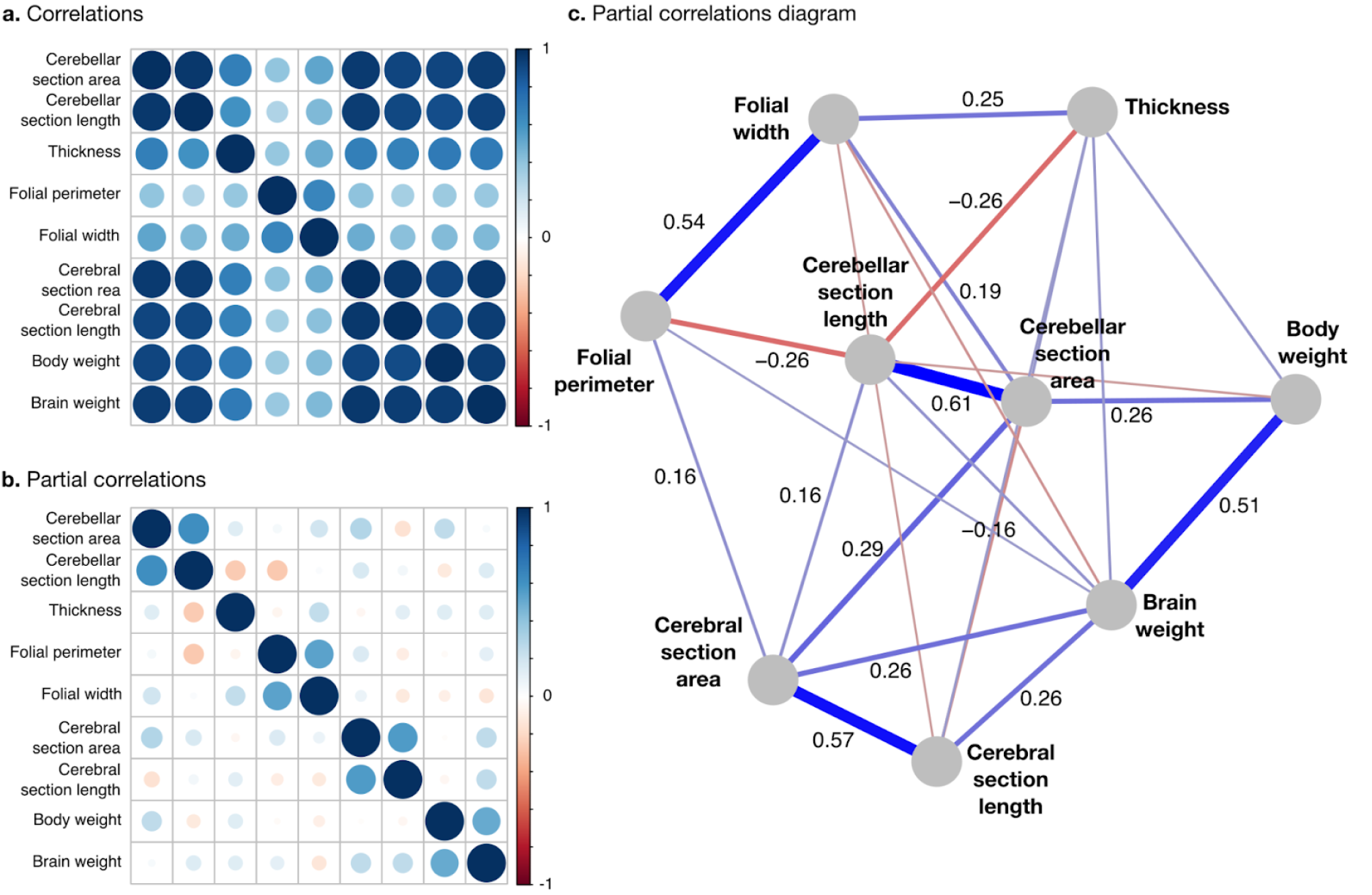
Correlation structure. **a.** Correlation matrix among all phenotypes (all values log10 transformed). **b.** Partial correlation matrix. **c.** Graph representation of the strongest positive (red) and negative (blue) partial correlations. All correlations are conditional to phylogenetic tree data.

Partial correlations (represented in a matrix in Fig. 7b and diagrammatically in Fig. 7c) were positive and strong between body weight and brain weight, cerebral section area and length, cerebellar section area and length, and folial width and perimeter. The most important positive partial correlation between cerebellum and cerebrum was between cerebral section area (a slice of cerebral volume), and cerebellar section area (a slice of cerebellar volume).

We also observed a positive partial correlation between the width of folia and the thickness of the molecular layer, which could be explained from the perspective of mechanical models of folding: thicker cortices are more difficult to fold, leading to wider folds. The 2 largest negative partial correlations were between cerebellar length and thickness of the molecular layer, and between cerebellar length and folial perimeter, indicating that larger cerebella tend to have relatively smaller, thinner folia than what could be expected for their size.

The phylogenetic principal component analysis of all cerebellar and cerebral measurements produced a 1^st^ principal component (PC1) which captured the largest part of the variance: 96.4% (Fig. 8). PC1 described a strong concerted change in body size and brain size, cerebellar and cerebral section area and length (size and folding), and less strong changes in folial width, folial perimeter and molecular layer thickness. PC1 describes the most important pattern of coordinated allometric variation (Jolicoeur 1963). In addition to allometric slopes computed from bivariate analyses, we can use the loadings of PC1 to compute a multivariate allometric scaling slope. This is obtained for any pair of variables as the ratio of their PC1 loadings. We will describe the allometric relationships this implies in the next sub-section.

**Figure 8.**
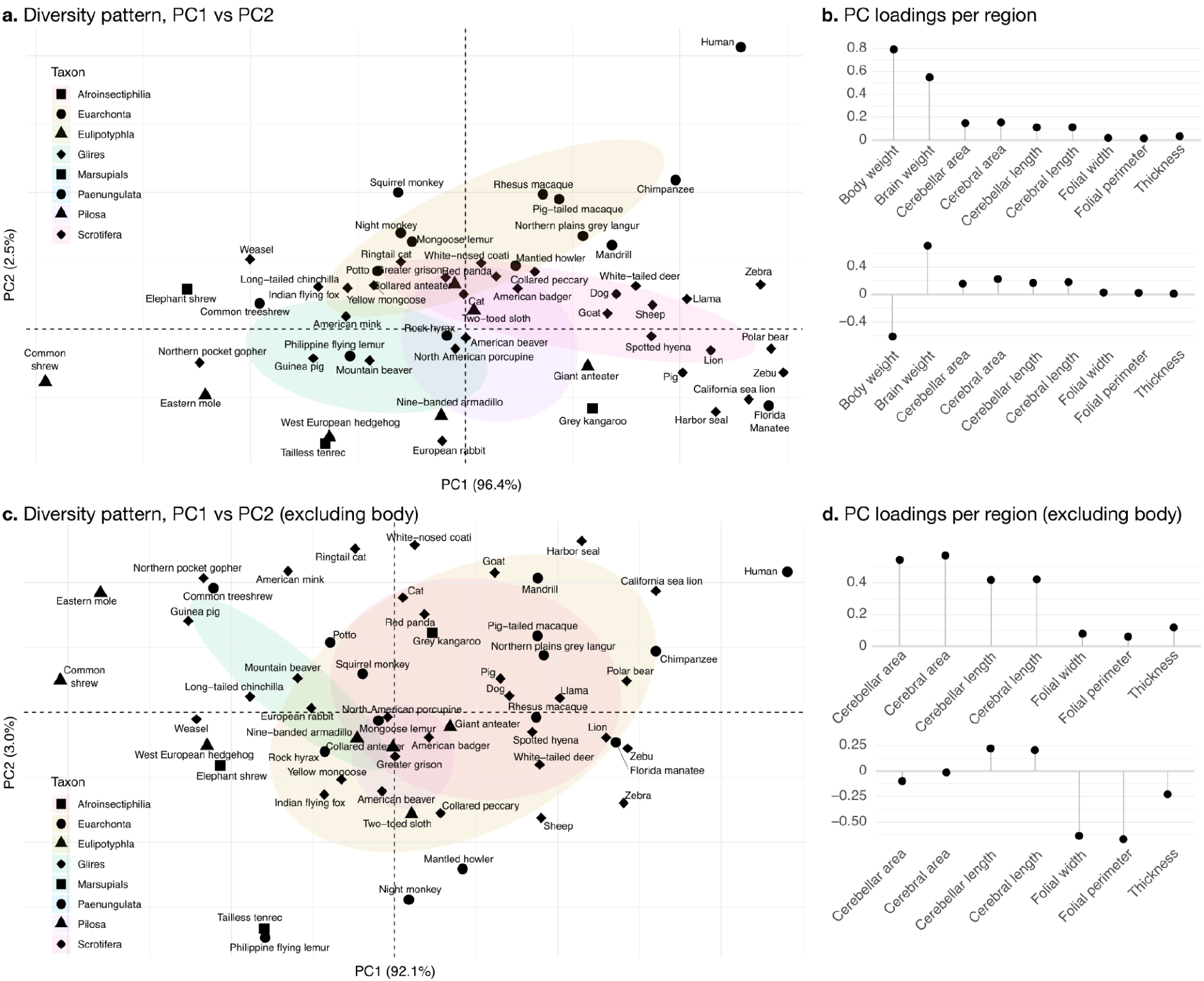
Principal components analysis. Primates, and especially humans, show particularly large brains given their body size. **a.** Pattern of neuroanatomical diversity (PC1 vs. PC2), including body weight. **b.** Loadings of the PC1 and PC2 axes displayed in panel a. **c.** Pattern of neuroanatomical diversity (PC1 vs. PC2), excluding body weight. **d.** Loadings of the PC1 and PC2 axes displayed in panel c.

PC2 conveys the main way in which individuals deviate from the dominant allometry pattern. The proportion of variance captured by PC2 was 2.5%. It mostly allowed uncoupling body size from changes in brain anatomy. Humans were here an important exception, with a small body size relative to brain size. To some extent, this was also the case for other primate species in our sample (chimpanzee, rhesus monkey, among others).

To better understand the patterns of neuroanatomical variation we performed an additional principal component analysis without including body size or brain size. This time PC1 captured 92.1% of the variance while PC2 captured 3.0%. The allometric pattern described by PC1 was similar to the previous one, with strong concerted changes in cerebellar and cerebral section area and length, and less strong changes in folial width, folial perimeter and molecular layer thickness. PC2 was different, and allowed to uncouple changes in folial width and folial perimeter from changes in molecular layer thickness. Here humans appeared among the species having the highest frequency of cerebellar folia (smaller folial width and perimeter).

### Ancestral character state estimation: Size and folding of the cerebrum and cerebellum increase and decrease concertedly through evolution

We computed ancestral estimations for all our phenotypes (Fig. 9). The results are well summarised by the ancestral estimations of the 1^st^ and 2^nd^ principal components. Fig. 9a and 9b show the ancestral estimation of PC1 and PC2 when including body size and brain size, while the analysis in Figs. 9c and 9d excludes them (to focus on neuroanatomy).

**Figure 9.**
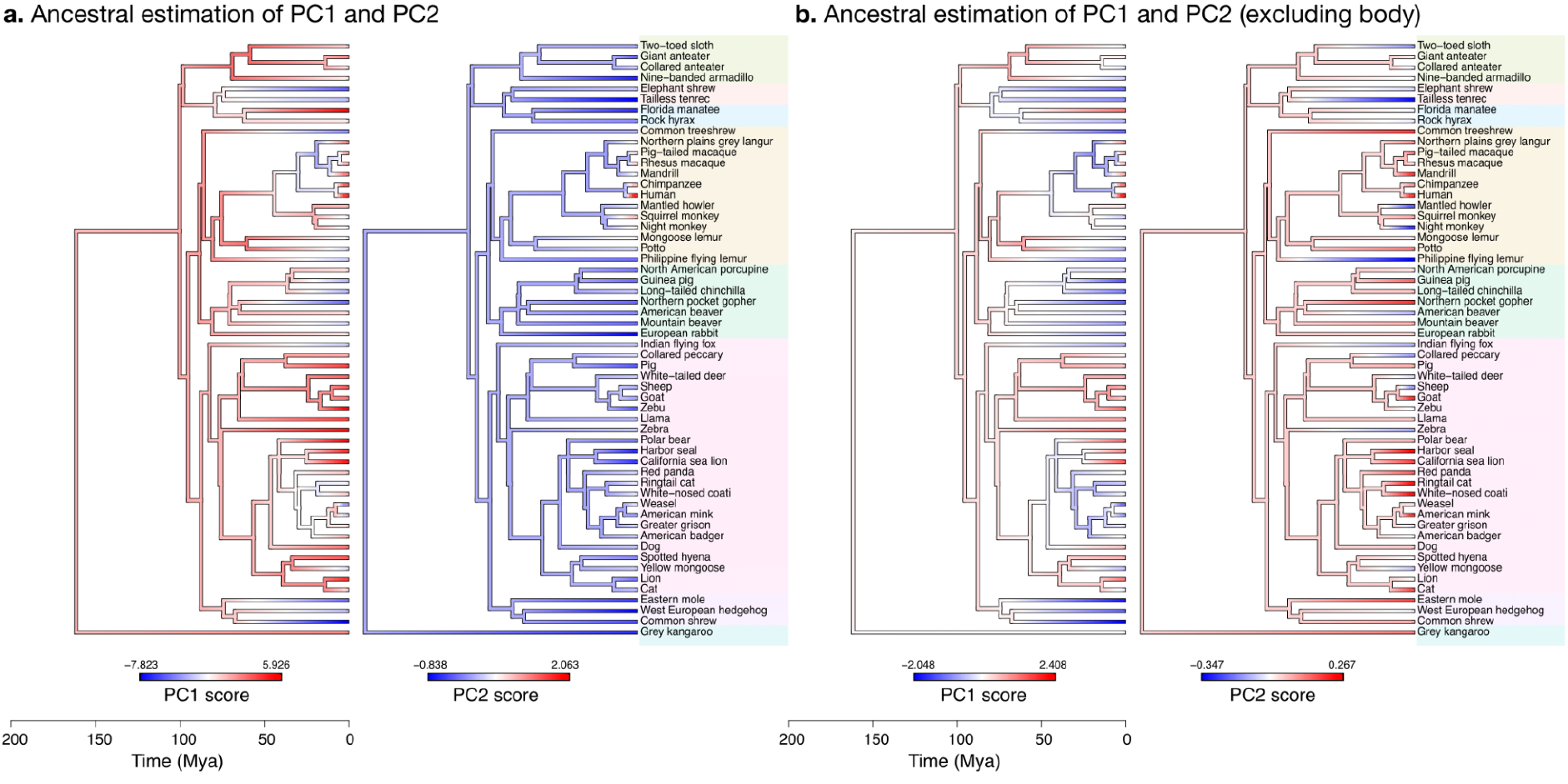
Estimation of ancestral neuroanatomical diversity patterns. Our analyses show a concerted change in body and brain size, and specific increase in cerebral and cerebellar volume in primates concomitant with an increased number of smaller folia. **a.** Ancestral estimation of PC1 and PC2 for neuroanatomical variables plus brain volume. PC1 captures 96.4% of the variance, PC2 captures 2.4%. **b.** Ancestral estimation of PC1 and PC2 for neuroanatomical variables, without brain volume. PC1 captures 91.4% of phenotypic variance, PC2 captures 3.6%.

The global pattern for PC1 in both analyses (Fig. 9a and 9b) were similar, and revealed the concomitant change in body size and brain size. Excluding those values, Fig. 9b highlights the specific increase in cerebral and cerebellar volume in primates.

The ancestral estimations for PC2 were more varied. When including body and brain size, they suggested that, within Euarchonta, primates have recently evolved brains which are larger than what could be expected from their body size. Excluding body and brain size (Fig. 9b) allowed us to better distinguish variation in the cerebrum and the cerebellum. We observed that cerebellar folia tended to be smaller in catarrhine primates (humans, macaques, etc.) but not in platyrrhine primates (squirrel monkey, night monkey, etc.). Although similar decreases in folial width and perimeter were present in other mammals with large brains such as the harbour seal, the combination of smaller folia and smaller body size (relative to brain size) seems to be characteristic of primates, which may have evolved a particularly folded cerebellum. The common ancestor of mammals is predicted to have had a folded cerebrum with a correspondingly folded cerebellum, similar to that of an American beaver.

### Allometry

Allometry is the study of the relationship between size and shape. When objects change only in volume without changing shape (isometric scaling), the length, surface area and volume of their parts change in a characteristic way: length as the square root of surface area, surface area as the 2/3 power of the volume, etc. We observed, for both cerebellum and cerebrum, that length scaled almost linearly with section area (instead of as the square root), indicating that cerebella are increasingly folded when larger (hyper-allometric scaling). Folial width, folial perimeter and molecular layer thickness are all measurements of length which should also scale as the square root of cerebellar section area in case of isometry. We observed a much smaller scaling factor, indicating that when comparing small and large brains, those measurements increase substantially less than what could be expected from increases in cerebellar size (hypo-allometric scaling).

#### Cerebellar folding increases with cerebellar size

The bivariate analyses (Fig. 10) zoom into the allometry of cerebellar and cerebral folding. In both cases, our measurement of length – a slice of the cerebellar and cerebral surface – correlated very strongly with the corresponding measurement of section area – a slice of cerebellar and cerebral volume. The increase of folding with size followed a similar law for both structures, with a scaling slope of 0.76–0.77. The scaling slopes were statistically significantly larger than 0.5, showing substantial hyper-allometry. The multivariate allometric slopes were similar to the bivariate slopes: 0.76 for the cerebellum and 0.73 for the cerebrum. In the case of the cerebrum, we confirmed that the Florida manatee was exceptionally unfolded: 3 times less cortical length than expected. For the cerebellum, we observed that humans had a particularly folded cerebellum: 1.5 times more cerebellar length than expected.

**Figure 10.**
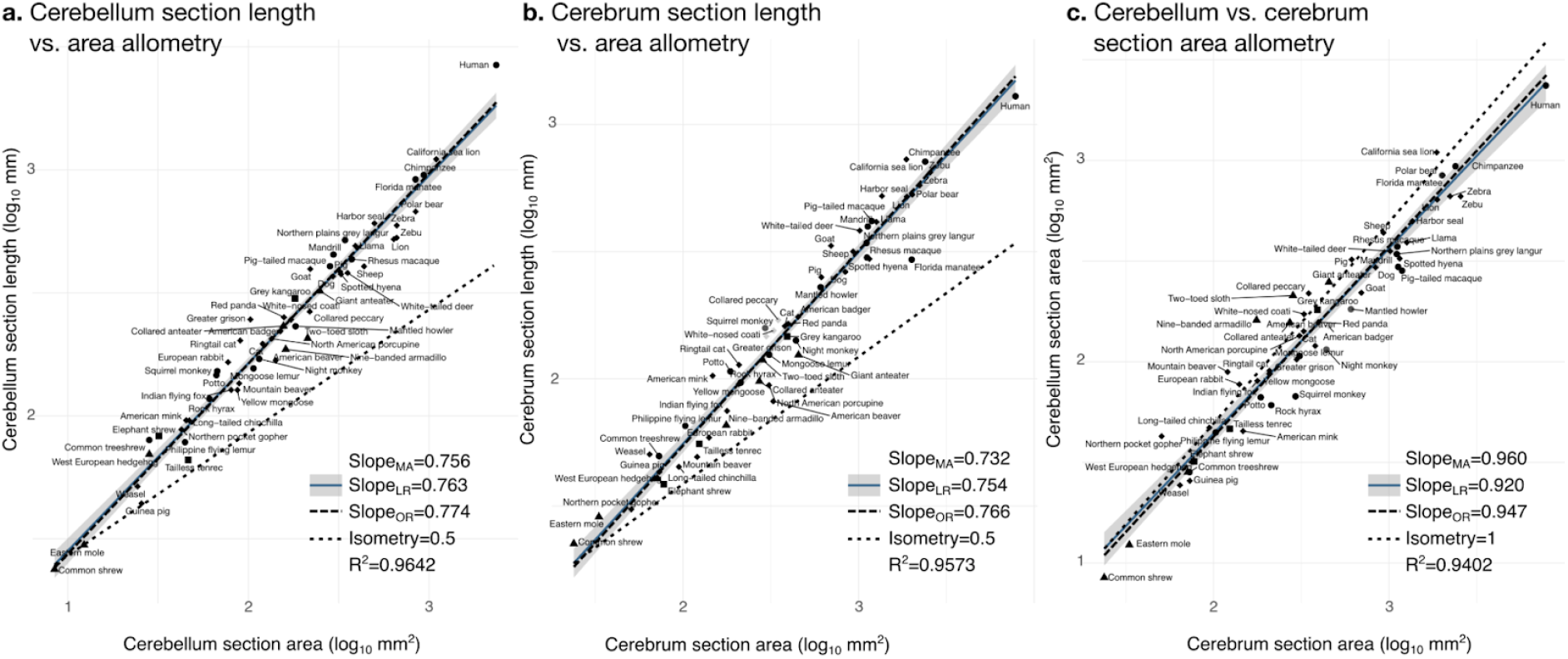
Cerebellum and cerebrum folding and allometry. The cerebellar and cerebral cortices are disproportionately larger than their volumes, as shown by their hyper-allometry. The cerebellum is slightly but statistically significantly hypo-allometric compared to the cerebrum **a.** Allometry of cerebellum section length vs. cerebellum section area. **b.** Allometry of cerebrum length vs. cerebrum section area. **c.** Allometry of cerebellum section area vs. cerebrum section area. MA: Multivariate allometry. LR: Linear regression with 95% confidence interval in grey. OR: Orthogonal regression.

The bivariate plot of cerebellum versus cerebral section area (Fig. 10c) showed a very strong correlation and suggests that the cerebellum is slightly proportionally smaller in species with large brains. In the case of isometry, the scaling slope should be 1. The observed scaling slope was slightly smaller: 0.96 when estimated through multivariate allometry, 0.95 when estimated through orthogonal regression, and 0.92 when estimated through linear regression. Although very close to 1, the difference is statistically significant: the isometric slope is outside the 95% confidence interval, suggesting a slight hypo-allometry of the cerebellum compared with the cerebrum.

#### Folial width, folial perimeter and the thickness of the molecular layer increase slightly with cerebellar size

The width and perimeter of cerebellar folia increased only slightly with cerebellar size (Fig. 11). The scaling was markedly hypo-allometric: 0.10 for folial perimeter and 0.14 for folial width where isometric scaling should be 0.5. The scaling of the molecular layer’s thickness was also hypo-allometric, but with a slightly higher scaling slope of 0.2. Folial width and perimeter appeared to be more variable than molecular layer thickness: linear regression with cerebellar section area captured only 28% of folial width variance, and 14% of folial perimeter variance, while it captured 64% of molecular layer thickness variance (Fig. 11c).

**Figure 11.**
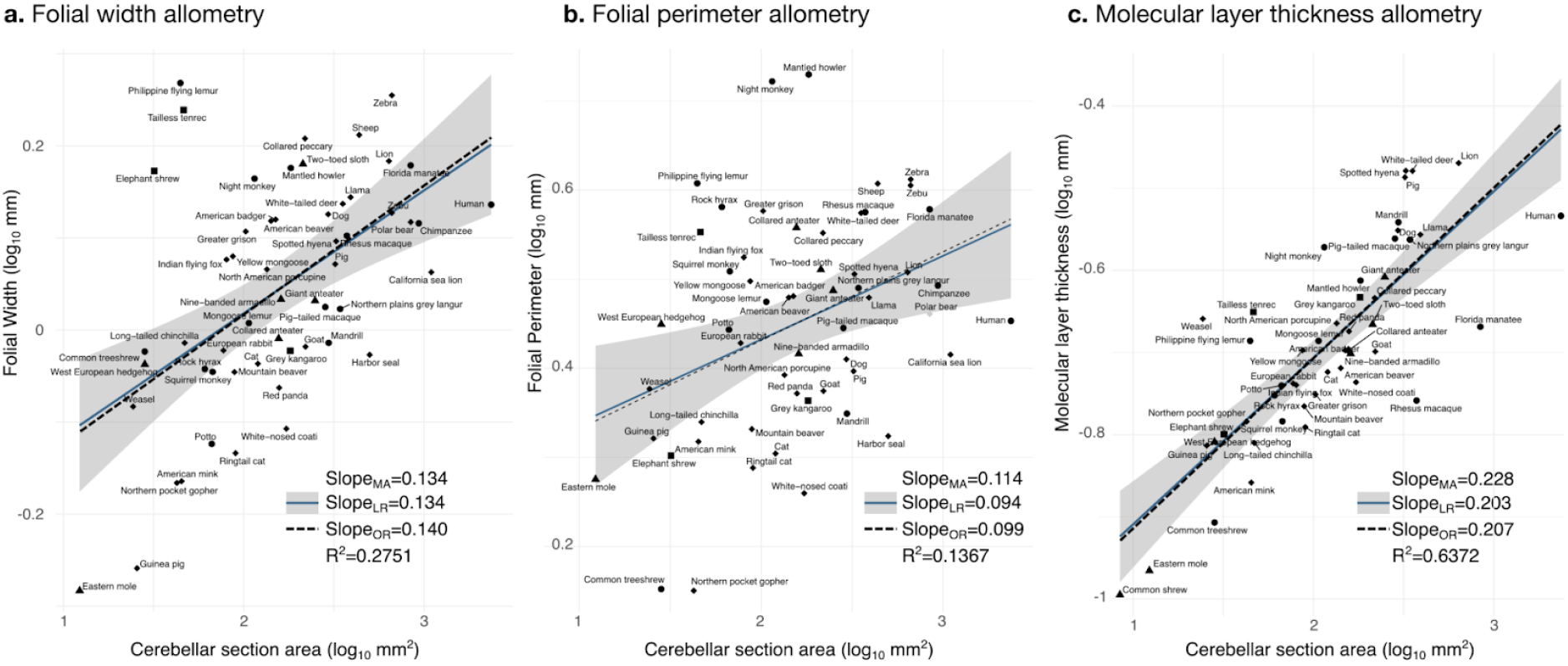
Allometry of folial width, perimeter and molecular layer thickness. The geometry of cerebellar folia and molecular layer thickness were largely conserved when compared with changes in total cerebellar size, as revealed by the small allometric slopes. **a.** Allometry of folial width vs. cerebellar section area (2-tailed p<1e-5). **b.** Allometry of folial perimeter vs. cerebellar section area (2-tailed p=0.005). **c.** Allometry of the thickness of the molecular layer vs. cerebellar section area (2-tailed p<1e-12). MA: Multivariate allometry. LR: Linear regression with 95% confidence interval in grey. OR: Orthogonal regression.

The mean thickness (t) of our molecular layer estimations across species was of t~200 μm, and the mean folial width (λ) across species was of λ~1 mm. The ratio of folial width (the wavelength of cerebellar folding) over molecular layer thickness is ~5, which is not far from the expected ratio for a model of mechanical wrinkling: λ = 2πt(μ/(3μ_s_))^1/3^, which approximates λ/t ~ 4.37 when the stiffness of the internal and external layers (μ_s_ and μ respectively) is similar (μ_s_~μ, see Tallinen et al. 2014).

## Discussion

> *Despite the impressive beauty of its wiring diagrams, the “neuronal machine” concept of the cerebellum remained vaguely defined as “a relatively simple machine devoted to some essential information processing.” I was frustrated enough at the Salishan meeting to ask what else experimentalists would need to uncover before we would be able to understand the meaning of these wiring diagrams. Someone equally frustrated replied that the available diagrams were too simple to construct even a primitive radio, so more information was urgently needed before any meaningful model could be conceived*.
>
> — M. Ito, *The Cerebellum* (2011)

Whereas the circuits of the cerebral cortex are characterised by the presence of profuse re-entering loops (most connections of the cerebral cortex originate in other regions of the same cerebral cortex), the cerebellum exhibits an almost perfect feedforward structure, with an organisation whose regularity and simplicity has baffled generations of experimentalists and theoreticians.

The impressive multi-scale regularity of cerebellar structure led to the idea of the “neuronal machine” (Eccles et al. 1967). Our quote from Ito (2011) illustrates the challenge of imagining how the obstinate repetition of a single interconnection pattern among a reduced number of components could lead to the complex cerebellar function. However, its impressive evolutionary conservation suggests that its function has to fulfil an important role, for motion and cognition (Whiting and Barton 2003, Barton and Venditti 2014, Magielse et al. 2022).

One possibility is that the complexity of cerebellar function should not be found at the level of the individual circuits – as would be the case for radio – but that those circuits would provide the substrate over which complexity would emerge, as waves in the ocean or patterns in a vibrating Chladni plate. A deeper understanding of the mesoscopic and macroscopic scales of organisation would be probably more appropriate to understanding phenomena unique to this level of organisation, complementing the extensive exploration of microscopic circuits and molecular properties of cerebellar cells.

Our results provide a closer look into the nature of macroscopic cerebellar anatomy, its diversity across mammalian species, and its evolution. We confirm the strong correlation between cerebral and cerebellar volumes across species (Herculano-Houzel 2010, Smaers et al. 2021), and we extended these results to show that the same strong relationship holds for cerebellar folding: larger cerebella appear to be disproportionately more folded than smaller ones.

Comparing the allometric relationships between section length and area suggests that a similar process lies behind the folding of the cerebrum and the cerebellum. The folding of the cerebral cortex has been the focus of intense research, both from the perspective of neurobiology and physics. Current models suggest that cortical folding should result from a buckling instability triggered by the growth of the cortical grey matter on top of the white matter core. In such systems, the growing layer should first expand without folding, increasing the stress in the core. But this configuration is unstable, and if growth continues stress is released through cortical folding. The wavelength of folding depends on cortical thickness, and folding models such as the one by Tallinen et al. (2014) predict a neocortical folding wavelength which corresponds well with the one observed in real cortices. Our results show that the same type of model can also predict the wavelength of cerebellar folding.

In the case of the cerebellum, it has been suggested that the external granular layer may play the role of the grey matter in the models of cerebral cortical folding (Lawton et al. 2019). During development, the external granular layer produces an astonishingly large number of small granule cells. These cells migrate past the Purkinje cells to constitute the granular layer (Leto et al. 2016). Alternatively, it would be possible that the expanding layer leading to cerebellar folding is the molecular layer, and that the growth is not driven by cellular proliferation, but by the growth of the dendritic arborization of Purkinje cells. Indeed, the growth of dendritic trees seems to be also an important factor in the growth of the cerebral cortex (Welker 1990). By the end of neuronal migration from the ventricular and outer subventricular zones, the cerebral cortex is still largely lissencephalic (Welker 1990). Most cerebral cortical folding starts after the end of neuronal migration and is concomitant with the development of cortico-cortical connectivity and the elaboration of neuronal dendritic trees.

We observed that folial width and folial perimeter changed only slightly compared with the large diversity in cerebellar size. This is similar to the observation that folding wavelength in the cerebral cortex across many primate species is relatively conserved (Heuer et al. 2019). In both cases, for the cerebellum and the cerebrum, the thickness of the expanding layer was also relatively conserved (compared with the much more important diversity of total cerebellar size), which could reflect the conservation of cerebellar circuits across species.

A striking characteristic of cerebellar anatomy is the multi-scale nature of its folding: in small cerebella we can observe only a first level of folding, but a pattern of folding within folding appears progressively as cerebellar size increases. In humans (see, for example, Fig. 1), we can distinguish at least 3 such levels of folding. The addition of new levels of folding could be the reason why we observe relatively smaller folial width in larger cerebella (Fig. 7c) such as those of humans, as new smaller folds develop on top of the older ones. In the human cerebrum, folding has been reported to show 1 level of folds on top of folds, or “frequency doubling” (Germanaud et al. 2012). This phenomenon has also been observed in swelling gels and mechanical models of folding (Mora and Boudaoud 2006). We could expect that such mechanical models should be able to produce additional levels of folding if more growth were allowed or if cortices were made thinner.

At the moment, we can only speculate about the role that the regular, strongly conserved, patterns we observe in cerebellar neuroanatomy may play in its functional organisation. The addition of an increasing number of folial “modules” and the constitution of a nested hierarchy of super modules should induce a similar modularity of white matter connections. Cerebellar folding could then constrain the cerebellum to develop a regular, hierarchical pattern of variation in fibre length with an associated modular pattern in timing of neuronal spikes. Then, through synaptic plasticity, this could lead to a preferential potentiation of neurones within the same folium, and next within their super-folium, etc. The timing of the development of cerebellar folding could also have an influence on the constitution of cerebellar networks, reinforcing the influence of early modules (trees) on late modules (leafs).

A finer understanding of the mesoscopic and macroscopic cerebellar neuroanatomical patterns would require more precise surface reconstructions in large numbers of species. The full reconstruction of the cerebellar surface is still challenging: to our knowledge, only 3 cerebella have been precisely reconstructed: 2 humans and 1 macaque (Sereno et al. 2020, Zheng et al. 2022). These two humans had markedly different cerebellar surface areas, one being >20% larger than the other, which highlight the necessity for increasing not only inter-individual precision, but also sample size. The previous work of Sultan and Braitenberg (1993) is remarkable in this respect, providing beautiful representations of the unfolded cerebellar surface for 15 different species. However, their method produced smaller cerebellar surface areas than in the same species using a more precise method, suggesting that it underestimates the actual values (Sereno et al. 2020, Zheng et al. 2022), and hence the magnitude of cerebellar folding.

Our analyses provide a different approach to the problem by quantifying precisely the folding of a single slice, allowing us to examine a much larger sample of 56 species. Our data is publicly available through MicroDraw, which enables researchers to easily reuse it to extend our analyses or to perform new ones. The methods we have implemented for the analysis of histological cerebellar data could be easily adapted to study some other brain structures, and it is our hope that they will facilitate the integration of a more rich comparative dimension to neuroimaging analyses.

## Acknowledgement

We thank Carol MacLeod, Fahad Sultan, David DiGregorio and Maria Castelló for discussions on cerebellar evolution and cerebellar folding, and Liam Revell and Julien Clavel for their help and advice with phylogenetic analyses. Funded by project NeuroWebLab (ANR-19-DATA-0025), DMOBE (ANR-21-CE45-0016), the European Union’s Horizon 2020 research and innovation programme under the Marie Skłodowska-Curie grant agreement No101033485 (KH Individual Fellowship), and the STIC-AmSud programme project STIC-AMSUD + CLANN 22-STIC-03.

---

**Supplemental table S1.** List of species included in this study, sorted alphabetically by orders and binomial name. Excluded_W&P_: excluded from width and period analyses. Excluded_Th_: excluded from analyses of molecular layer thickness.

| **Binomial name** | **English name** | **Provenance** | **Excluded<sub>W&P</sub>** | **Excluded<sub>Th</sub>** |
| --- | --- | --- | --- | --- |
| **Afroinsectiphilia** |  |  |  |  |
| *Tenrec ecaudatus* | Tailless tenrec | Brainmuseum |  |  |
| *Elephantulus myurus* | Elephant shrew | Brainmuseum |  |  |
| **Euarchonta** |  |  |  |  |
| *Alouatta palliata* | Mantled howler | Brainmuseum |  |  |
| *Aotus trivirgatus* | Night monkey | Brainmuseum |  |  |
| *Cynocephalus volans* | Philippine flying lemur | Brainmuseum |  |  |
| *Eulemur mongoz* | Mongoose lemur | Brainmuseum |  |  |
| *Homo sapiens* | Human | BigBrain Project |  |  |
| *Macaca mulatta* | Rhesus macaque | Brainmaps |  |  |
| *Macaca nemestrina* | Pig-tailed macaque | Brainmuseum |  |  |
| *Mandrillus sphinx* | Mandrill | Brainmuseum |  |  |
| *Pan troglodytes* | Chimpanzee | Brainmuseum |  | Yes |
| *Perodicticus potto* | Potto | Brainmuseum |  |  |
| *Saimiri sciureus* | Squirrel monkey | Brainmuseum |  |  |
| *Semnopithecus entellus* | Northern plains grey langur | Brainmuseum |  |  |
| *Tupaia glis* | Common tree shrew | Brainmuseum |  |  |
| **Eulipotyphla** |  |  |  |  |
| *Choloepus didactylus* | Two-toed sloth | Brainmuseum |  |  |
| *Dasypus novemcinctus* | Nine-banded armadillo | Brainmuseum |  |  |
| *Myrmecophaga tridactyla* | Giant anteater | Brainmuseum |  |  |
| *Tamandua tetradactyla* | Collared anteater | Brainmuseum |  |  |
| **Glires** |  |  |  |  |
| *Aplodontia rufa* | Mountain beaver | Brainmuseum |  |  |
| *Castor canadensis* | American beaver | Brainmuseum |  |  |
| *Cavia porcellus* | Guinea pig | Brainmuseum |  |  |
| *Chinchilla lanigera* | Long-tailed chinchilla | Brainmuseum |  |  |
| *Erethizon dorsatum* | North American porcupine | Brainmuseum |  |  |
| *Oryctolagus cuniculus* | European rabbit | Brainmuseum |  |  |
| *Thomomys talpoides* | Northern pocket gopher | Brainmuseum |  |  |
| **Marsupials** |  |  |  |  |
| *Macropus fuliginosus* | Grey kangaroo | Brainmuseum |  |  |
| **Paenungulata** |  |  |  |  |
| *Procavia capensis* | Rock hyrax | Brainmuseum |  |  |
| *Trichechus manatus latirostris* | Florida manatee | Brainmuseum |  |  |
| **Pilosa** |  |  |  |  |
| *Erinaceus europaeus* | West European hedgehog | Brainmuseum |  |  |
| *Scalopus aquaticus* | Eastern mole | Brainmuseum |  |  |
| *Sorex araneus* | Common shrew | Brainmuseum | Yes |  |
| **Scrotifera** |  |  |  |  |
| *Bos taurus indicus* | Zebu | Brainmuseum |  | Yes |
| *Capra hircus domestica* | Goat | Brainmuseum |  |  |
| *Lama glama* | Llama | Brainmuseum |  |  |
| *Odocoileus virginianus* | White-tailed deer | Brainmuseum |  |  |
| *Ovis aries domestica* | Sheep | Brainmuseum |  | Yes |
| *Pecari tajacu* | Collared peccary | Brainmuseum |  |  |
| *Sus scrofa domesticus* | Pig | Brainmuseum |  |  |
| *Ailurus fulgens* | Red panda | Brainmuseum |  |  |
| *Bassariscus astutus* | Ringtail | Brainmuseum |  |  |
| *Canis familiaris* | Dog | Brainmuseum |  |  |
| *Crocuta crocuta* | Spotted hyena | Brainmuseum |  |  |
| *Cynictis penicillata* | Yellow mongoose | Brainmuseum |  |  |
| *Felis catus* | Cat | Brainmuseum |  |  |
| *Galictis vittata* | Greater grison | Brainmuseum |  |  |
| *Mustela erminea* | Weasel | Brainmuseum |  |  |
| *Nasua narica* | White-nosed coati | Brainmuseum |  |  |
| *Neovison vison* | American mink | Brainmuseum |  |  |
| *Panthera leo* | Lion | Brainmuseum |  |  |
| *Taxidea taxus* | American badger | Brainmuseum |  |  |
| *Ursus maritimus* | Polar bear | Brainmuseum |  | Yes |
| *Pteropus giganteus* | Indian flying fox | Brainmuseum |  |  |
| *Equus burchellii* | Burchell's zebra | Brainmuseum |  | Yes |
| *Phoca vitulina* | Harbor seal | Brainmuseum |  | Yes |
| *Zalophus californianus* | California sea lion | Brainmuseum |  | Yes |

**Supplemental Figure Fig. 1.**
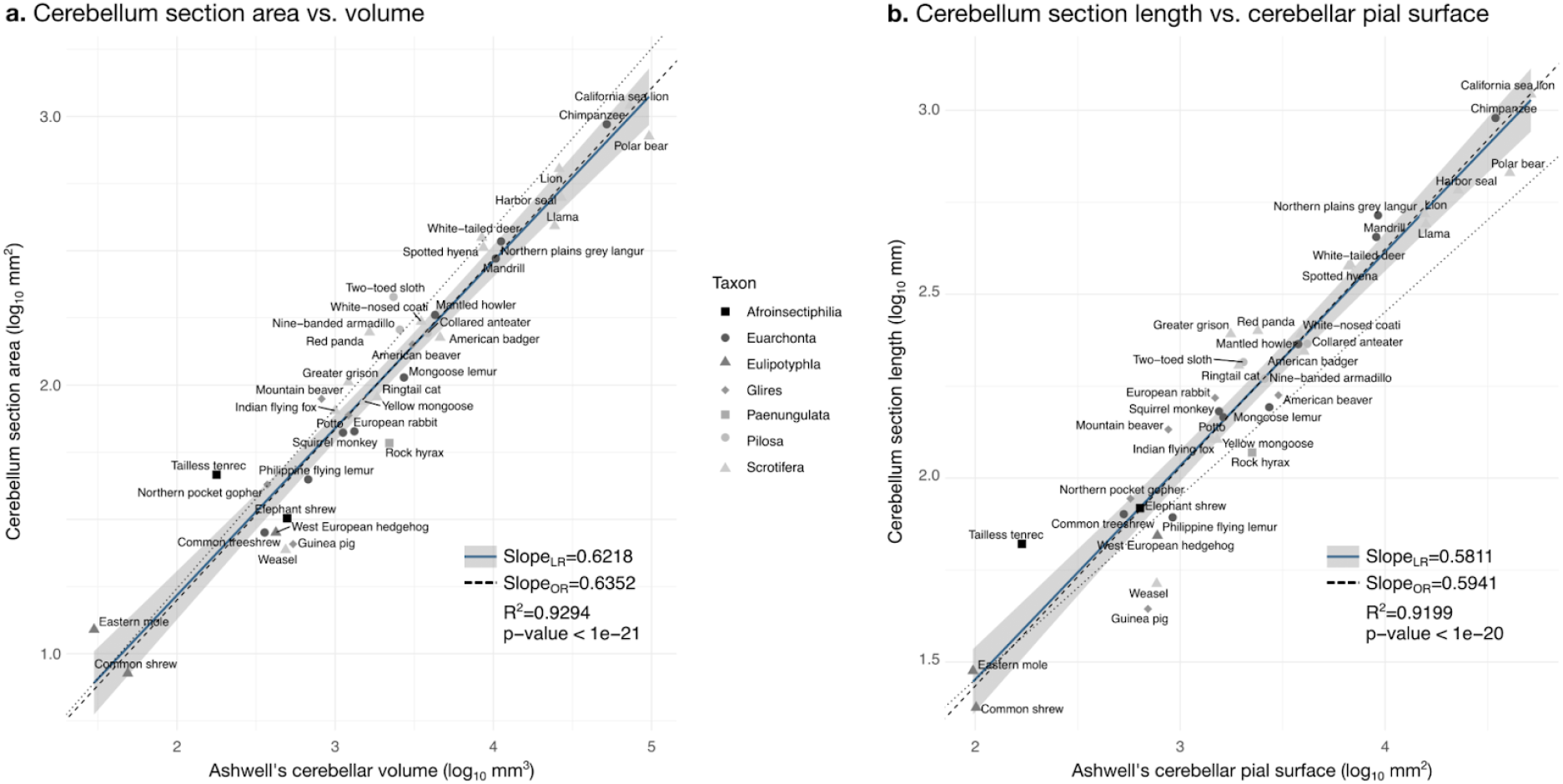
Correlation with measurements in Ashwell (2020). / Validation of neuroanatomical measurements. Comparison of our cerebellar section area measurement **(a)** and cerebellar section length **(b)** with cerebellar volume and surface measurements from Ashwell (2020). Measurements correlate significantly (2-tailed p<1e-20). LR: Linear regression with 95% confidence interval in grey. OR: Orthogonal regression.

## Notes

https://github.com/neuroanatomy/comp-cb-folding

https://microdraw.pasteur.fr/project/brainmuseum-cb

### Competing Interest Statement

The authors have declared no competing interest.

### Summary of Updates

- species names are now displayed in english in all figures - all model fitting is done with mvMorph, instead of using rphylopars for imputation as in the first version - 2 rhesus monkeys were displayed in Fig. 1, but only one of those was used.

## References

Amunts, K., Lepage, C., Borgeat, L., Mohlberg, H., Dickscheid, T., Rousseau, M.-É., Bludau, S., Bazin, P.-L., Lewis, L. B., Oros-Peusquens, A.-M., Shah, N. J., Lippert, T., Zilles, K., & Evans, A. C. (2013). BigBrain: An Ultrahigh-Resolution 3D Human Brain Model. Science (Vol. 340, Issue 6139, pp. 1472–1475). American Association for the Advancement of Science (AAAS). https://doi.org/10.1126/science.1235381

Ballarin, C., Povinelli, M., Granato, A., Panin, M., Corain, L., Peruffo, A., & Cozzi, B. (2016). The Brain of the Domestic Bos taurus: Weight, Encephalization and Cerebellar Quotients, and Comparison with Other Domestic and Wild Cetartiodactyla. In P. Raia (Ed.), PLOS ONE (Vol. 11, Issue 4, p. e0154580). Public Library of Science (PLoS). https://doi.org/10.1371/journal.pone.0154580

Barton, R. A., & Venditti, C. (2014). Rapid Evolution of the Cerebellum in Humans and Other Great Apes. Current Biology (Vol. 24, Issue 20, pp. 2440–2444). Elsevier BV. https://doi.org/10.1016/j.cub.2014.08.056

Burger, J. R., George, M. A., Jr., Leadbetter, C., & Shaikh, F. (2019). The allometry of brain size i n mammals. In Journal of Mammalogy (Vol. 100, Issue 2, pp. 276–283). Oxford University Press (OUP). https://doi.org/10.1093/jmammal/gyz043

Clavel, J., Escarguel, G., & Merceron, G. (2015). mvMORPH: an R package for fitting multivariate evolutionary models to morphometric data. Methods in Ecology and Evolution (Vol. 6, Issue 11, pp. 1311–1319). Wiley. https://doi.org/10.1111/2041-210x.12420

Eccles, J. C., Ito, M., & Szentágothai, J. (1967). The cerebellum as a neuronal machine. Springer-Verlag. https://doi.org/10.1007/978-3-662-13147-3

Foubet, O. Trejo, M., & Toro, R. (2019). Mechanical morphogenesis and the development of neocortical organisation. Cortex, 118, 315–326. https://doi.org/10.1016/j.cortex.2018.03.005

Franze, K. (2013). The mechanical control of nervous system development. Development, 140(15), 3069–3077. https://doi.org/10.1242/dev.079145

Germanaud, D., Lefèvre, J., Toro, R., Fischer, C., Dubois, J., Hertz-Pannier, L., & Mangin, J.-F. (2012). Larger is twistier: Spectral analysis of gyrification (SPANGY) applied to adult brain size polymorphism. NeuroImage (Vol. 63, Issue 3, pp. 1257–1272). https://doi.org/10.1016/j.neuroimage.2012.07.053

Herculano-Houzel. (2010). Coordinated scaling of cortical and cerebellar numbers of neurons. Frontiers in Neuroanatomy https://doi.org/10.3389/fnana.2010.00012

Heuer, K. and Toro, R. (2019). Role of mechanical morphogenesis in the development and evolution of the neocortex. Physics of Life Reviews, 31, 233–239. https://doi.org/10.1016/j.plrev.2019.01.012

Heuer, K., Gulban, O. F., Bazin, P.-L., Osoianu, A., Valabregue, R., Santin, M., Herbin, M., & Toro, R. (2019). Evolution of neocortical folding: A phylogenetic comparative analysis of MRI from 34 primate species. Cortex (Vol. 118, pp. 275–291). Elsevier BV. https://doi.org/10.1016/j.cortex.2019.04.011

Ito, Masao (2011). The Cerebellum. ISBN 0-13-705068-2. FT Press Science, 1st Edition.

Jiang, S., Guan, Y., Chen, S., Jia, X., Ni, H., Zhang, Y., Han, Y., Peng, X., Zhou, C., Li, A., Luo, Q., & Gong, H. (2020). Anatomically revealed morphological patterns of pyramidal neurons in layer 5 of the motor cortex. Scientific Reports (Vol. 10, Issue 1). Springer Science and Business Media LLC. https://doi.org/10.1038/s41598-020-64665-2

Jolicoeur, P. (1963). 193. The Multivariate Generalization of the Allometry Equation. Biometrics (Vol. 19, Issue 3, p. 497). JSTOR. https://doi.org/10.2307/2527939

Kaneko, M., Yamaguchi, K., Eiraku, M., Sato, M., Takata, N., Kiyohara, Y., Mishina, M., Hirase, H., Hashikawa, T., & Kengaku, M. (2011). Remodeling of Monoplanar Purkinje Cell Dendrites during Cerebellar Circuit Formation. PLoS ONE (Vol. 6, Issue 5, p. e20108). https://doi.org/10.1371/journal.pone.0020108

Kroenke, C. D., & Bayly, P. V. (2018). How Forces Fold the Cerebral Cortex. Journal of Neuroscience (Vol. 38, Issue 4, pp. 767–775). Society for Neuroscience. https://doi.org/10.1523/jneurosci.1105-17.2017

Kumar, S., Stecher, G., Suleski, M., & Hedges, S. B. (2017). TimeTree: A Resource for Timelines, Timetrees, and Divergence Times. Molecular Biology and Evolution (Vol. 34, Issue 7, pp. 1812–1819). Oxford University Press (OUP). https://doi.org/10.1093/molbev/msx116

Lawton, A. K., Engstrom, T., Rohrbach, D., Omura, M., Turnbull, D. H., Mamou, J., Zhang, T., Schwarz, J. M., & Joyner, A. L. (2019). Cerebellar folding is initiated by mechanical constraints on a fluid-like layer without a cellular pre-pattern. eLife (Vol. 8). eLife Sciences Publications, Ltd. https://doi.org/10.7554/elife.45019

Leto, K., Arancillo, M., Becker, E. B. E., Buffo, A., Chiang, C., Ding, B., Dobyns, W. B., Dusart, I., Haldipur, P., Hatten, M. E., Hoshino, M., Joyner, A. L., Kano, M., Kilpatrick, D. L., Koibuchi, N., Marino, S., Martinez, S., Millen, K. J., Millner, T. O., … Hawkes, R. (2015). Consensus Paper: Cerebellar Development. The Cerebellum (Vol. 15, Issue 6, pp. 789–828). https://doi.org/10.1007/s12311-015-0724-2

Llinares-Benadero, C., & Borrell, V. (2019). Deconstructing cortical folding: genetic, cellular and mechanical determinants. Nature Reviews Neuroscience (Vol. 20, Issue 3, pp. 161–176). Springer Science and Business Media LLC. https://doi.org/10.1038/s41583-018-0112-2

Magielse, N., Heuer, K., Toro, R., Schutter, D. J. L. G., & Valk, S. L. (2022). A Comparative Perspective on the Cerebello-Cerebral System and Its Link to Cognition. The Cerebellum. Springer Science and Business Media LLC. https://doi.org/10.1007/s12311-022-01495-0

Mora, T., & Boudaoud, A. (2006). Buckling of swelling gels. European Physical Journal E (Vol. 20, Issue 2, pp. 119–124). https://doi.org/10.1140/epje/i2005-10124-5

Mota, B., & Herculano-Houzel, S. (2015). Cortical folding scales universally with surface area and thickness, not number of neurons. Science (Vol. 349, Issue 6243, pp. 74–77). American Association for the Advancement of Science (AAAS). https://doi.org/10.1126/science.aaa9101

Revell, L. J. (2011). Phytools: an R package for phylogenetic comparative biology (and other things). Methods in Ecology and Evolution (Vol. 3, Issue 2, pp. 217–223). Wiley. https://doi.org/10.1111/j.2041-210x.2011.00169.x

Sereno, M. I., Diedrichsen, J., Tachrount, M., Testa-Silva, G., d’Arceuil, H., & De Zeeuw, C. (2020). The human cerebellum has almost 80% of the surface area of the neocortex. Proceedings of the National Academy of Sciences (Vol. 117, Issue 32, pp. 19538–19543). Proceedings of the National Academy of Sciences. https://doi.org/10.1073/pnas.2002896117

Shigeno, S., Andrews, P. L. R., Ponte, G., & Fiorito, G. (2018). Cephalopod Brains: An Overview of Current Knowledge to Facilitate Comparison With Vertebrates. Frontiers in Physiology (Vol. 9). Frontiers Media SA. https://doi.org/10.3389/fphys.2018.00952

Smaers, J. B., Turner, A. H., Gómez-Robles, A., & Sherwood, C. C. (2018). A cerebellar substrate for cognition evolved multiple times independently in mammals. eLife (Vol. 7). https://doi.org/10.7554/elife.35696

Smaers, J. B., Rothman, R. S., Hudson, D. R., Balanoff, A. M., Beatty, B., Dechmann, D. K. N., de Vries, D., Dunn, J. C., Fleagle, J. G., Gilbert, C. C., Goswami, A., Iwaniuk, A. N., Jungers, W. L., Kerney, M., Ksepka, D. T., Manger, P. R., Mongle, C. S., Rohlf, F. J., Smith, N. A., … Safi, K. (2021). The evolution of mammalian brain size. Science Advances (Vol. 7, Issue 18). American Association for the Advancement of Science (AAAS). https://doi.org/10.1126/sciadv.abe2101

Sultan F. & Braitenberg V. (1993) Shapes and sizes of different mammalian cerebella. A study in quantitative comparative neuroanatomy. J. Hirnforsch 34(1):79–92. PMID: 8376757.

Tallinen, T., Chung, J. Y., Biggins, J. S., & Mahadevan, L. (2014). Gyrification from constrained cortical expansion. Proceedings of the National Academy of Sciences, 111(35), 12667–12672. https://doi.org/10.1073/pnas.1406015111

Toro, R., & Burnod, Y. (2005). A Morphogenetic Model for the Development of Cortical Convolutions. Cerebral Cortex, 15(12), 1900–1913. https://doi.org/10.1093/cercor/bhi068

Toro, R. (2012). On the Possible Shapes of the Brain. Evolutionary Biology, 39(4), 600–612. https://doi.org/10.1007/s11692-012-9201-8

Van der Walt, S., Schönberger, J. L., Nunez-Iglesias, J., Boulogne, F., Warner, J. D., Yager, N., Gouillart, E., & Yu, T. (2014). scikit-image: image processing in Python. PeerJ (Vol. 2, p. e453). PeerJ. https://doi.org/10.7717/peerj.453

Van Essen, D. C. (2020). A 2020 view of tension-based cortical morphogenesis. In Proceedings of the National Academy of Sciences (Vol. 117, Issue 52, pp. 32868–32879). Proceedings of the National Academy of Sciences. https://doi.org/10.1073/pnas.2016830117

Welker, W. (1990). Why does cerebral cortex fissure and fold? Cerebral Cortex (pp. 3–136). Springer US. https://doi.org/10.1007/978-1-4615-3824-0_1

Whiting, B. A., & Barton, R. A. (2003). The evolution of the cortico-cerebellar complex in primates: anatomical connections predict patterns of correlated evolution. Journal of Human Evolution (Vol. 44, Issue 1, pp. 3–10). Elsevier BV. https://doi.org/10.1016/s0047-2484(02)00162-8

Zheng, J., Yang, Q., Makris, N., Huang, K., Liang, J., Ye, C., Yu, X., Tian, M., Ma, T., Mou, T., Guo, W., Kikinis, R., & Gao, Y. (2022). Three-Dimensional Digital Reconstruction of the Cerebellar Cortex: Lobule Thickness, Surface Area Measurements, and Layer Architecture. The Cerebellum. Springer Science and Business Media LLC. https://doi.org/10.1007/s12311-022-01390-8

